# Automating Mendelian randomization through machine learning to construct a putative causal map of the human phenome

**DOI:** 10.1101/173682

**Authors:** Gibran Hemani, Jack Bowden, Philip Haycock, Jie Zheng, Oliver Davis, Peter Flach, Tom Gaunt, George Davey Smith

**Author notes:** Equal contribution (Tom Gaunt, George Davey Smith).

## Abstract

A major application for genome-wide association studies (GWAS) has been the emerging field of causal inference using Mendelian randomization (MR), where the causal effect between a pair of traits can be estimated using only summary level data. MR depends on SNPs exhibiting vertical pleiotropy, where the SNP influences an outcome phenotype only through an exposure phenotype. Issues arise when this assumption is violated due to SNPs exhibiting horizontal pleiotropy. We demonstrate that across a range of pleiotropy models, instrument selection will be increasingly liable to selecting invalid instruments as GWAS sample sizes continue to grow. Methods have been developed in an attempt to protect MR from different patterns of horizontal pleiotropy, and here we have designed a mixture-of-experts machine learning framework (MR-MoE 1.0) that predicts the most appropriate model to use for any specific causal analysis, improving on both power and false discovery rates. Using the approach, we systematically estimated the causal effects amongst 2407 phenotypes. Almost 90% of causal estimates indicated some level of horizontal pleiotropy. The causal estimates are organised into a publicly available graph database (http://eve.mrbase.org), and we use it here to highlight the numerous challenges that remain in automated causal inference.

## Introduction

Mendelian randomization (MR) (1,2) exploits vertical pleiotropy to infer the causal relationships between phenotypes. Suppose that one trait (the exposure) causally influences another (the outcome). If a SNP influences the outcome through the exposure then the SNP is exhibiting vertical pleiotropy (see Box 1). Such a genetic variant is considered to be a valid instrumental variable if it only influences the outcome through the exposure (the exclusion restriction assumption). Vertical pleiotropy can be exploited to mimic a randomised controlled trial, enabling a causal estimate to be made by comparing the outcome phenotypes between those individuals that have the exposure-increasing allele against those who do not, although not without caveats (3). Multiple independent genetic variants for a particular exposure can be used jointly to improve causal inference, because a) each variant represents an independent natural experiment, and an overall causal estimate can be obtained by meta-analysing the single estimates from each instrument; and b) potential bias arising from the exclusion restriction assumption can be detected or corrected by evaluating the consistency of effects across instruments (4–9).

Genome-wide association studies (GWAS) have identified potential instrumental variables for thousands of phenotypes (10). Recent developments in MR have enabled knowledge of instrumental variables to be applied using either individual-level or only summary-level data (known as two-sample MR, 2SMR) (11). Here, in order to infer the causal effect of an exposure on an outcome all that is required are the genetic effects of the instrumenting SNP on the exposure and the outcome. This has three major advantages. First, GWAS summary data are non-disclosive and often publicly available. Second, causal inference can be made between phenotypes even if they have not been measured in the same samples, limiting possible MR analyses only by the availability of GWAS summary data for the traits in question (12). Third, because each instrumental variable mimics an independent randomization study, we can view the causal inference problem through the simple and widely understood prism of a meta-analysis (6).

Problems with obtaining unbiased causal effects can arise, however, if the genetic instruments exhibit horizontal pleiotropy, where they influence the outcome through a pathway other than the exposure (Box 1). The extent of this phenomenon is not to be understated, and many methods have been developed that attempt to reliably obtain unbiased causal estimates under specific models of horizontal pleiotropy (4–7,9,13). It is considered best practice to triangulate estimates from a range of MR methods (and other experimental designs) when presenting causal estimates so as to assess the potential for different horizontal pleiotropy models and other violations of MR assumptions (14). However there are circumstances in which it is desirable to consolidate across many methods to obtain a single MR estimate. First, it could be of interest to screen links between thousands of traits for being potential causal associations, in which case a critical evaluation of each causal estimate of interest may not be possible or convenient. Second, though the simplest method, based on inverse-variance weighted (IVW) meta analysis (11), is most statistically powerful under no horizontal pleiotropy, it can have high false positive or low true positive rates in the presence of horizontal pleiotropy compared to other methods. Pleiotropy has been hypothesised to be universal (15), though the degree and nature (e.g. horizontal versus vertical, see Box 1) to which this may be the case is contested (16,17), hence defaulting to the IVW method in the first instance and using other methods as sensitivity analyses may not be appropriate. Third, if different methods disagree it is not possible to know which is correct because the true nature of horizontal pleiotropy exhibited by the instruments is not known. Fourth, the available methods do not cover all possible models of horizontal pleiotropy, and therefore an automated method for instrument selection may be necessary.

In this paper we introduce two innovations towards improving the reliability of MR estimates. First, we present an approach to discard genetic variants that are likely to be invalid. Second, we hypothesised that characteristics of the summary data could indicate which method would be most reliable, and we introduce new machine learning approaches that attempt to automate both instrument and method selection. Using curated GWAS summary data for thousands of phenotypes (12), we use these new methods to construct a graph of millions of causal estimates. Motivated by the recent avalanche (18) of 2SMR publications, with similar (19,20) or contradicting (21,22) conclusions despite using the same data (18), we developed a graph database to represent these estimates in a consistent manner. We consider this to be a ‘working draft’ of the causal map of the human phenome, but raise caution throughout that its interpretation is far from straightforward, and that future corrections and refinements will inevitably follow as data grows and 2SMR methods evolve.

#### Box 1: Pleiotropic mechanisms

Pleiotropy is the phenomenon whereby a single genetic variant influences multiple phenotypes. The mechanisms behind it can arise through many different mechanisms, for example a single variant influences multiple genes; a single gene generates multiple gene products; a single gene product has multiple cellular functions; or a single cellular function influences multiple phenotypes (23). How we classify the pleiotropic mechanism depends on question and context, and in this paper we discuss pleiotropy in two forms - vertical and horizontal. Vertical pleiotropy has also been known as mediated pleiotropy (24), because the genetic variant influences one trait, which in turn influences a second trait. It has also been termed, in some contexts, Type II pleiotropy (16) and secondary pleiotropy (25). It is this form of pleiotropy that MR assumes to make causal inference between two traits.

By contrast, horizontal pleiotropy can manifest in a myriad of different ways, but the basic principle is that the genetic variant influences two different phenotypes through independent pathways. Depending on context, horizontal pleiotropy can also be termed biological pleiotropy (24), Type I pleiotropy (16), developmental pleiotropy or selectional pleiotropy (26). From the perspective of MR, horizontal pleiotropy arises when a SNP influences an outcome trait through some pathway other than the exposure trait.

A genetic variant can exhibit both horizontal and vertical pleiotropy in different MR analyses. For example, consider a scenario where a SNP influences the expression levels of two genes (X and Y), and gene X influences trait A while gene Y influences trait B. Then:

- The genetic variant is exhibiting a vertical pleiotropic effect on trait A and the expression level of gene X. Here, all other assumptions being met, the genetic variant would be a valid instrument if used to estimate the causal effect of the gene expression level on trait A. However,
- the SNP is exhibiting a horizontal pleiotropic effect on trait A and trait B through the two gene expression levels. Here the genetic variant would be invalid if used to estimate the causal effect of trait A on trait B.

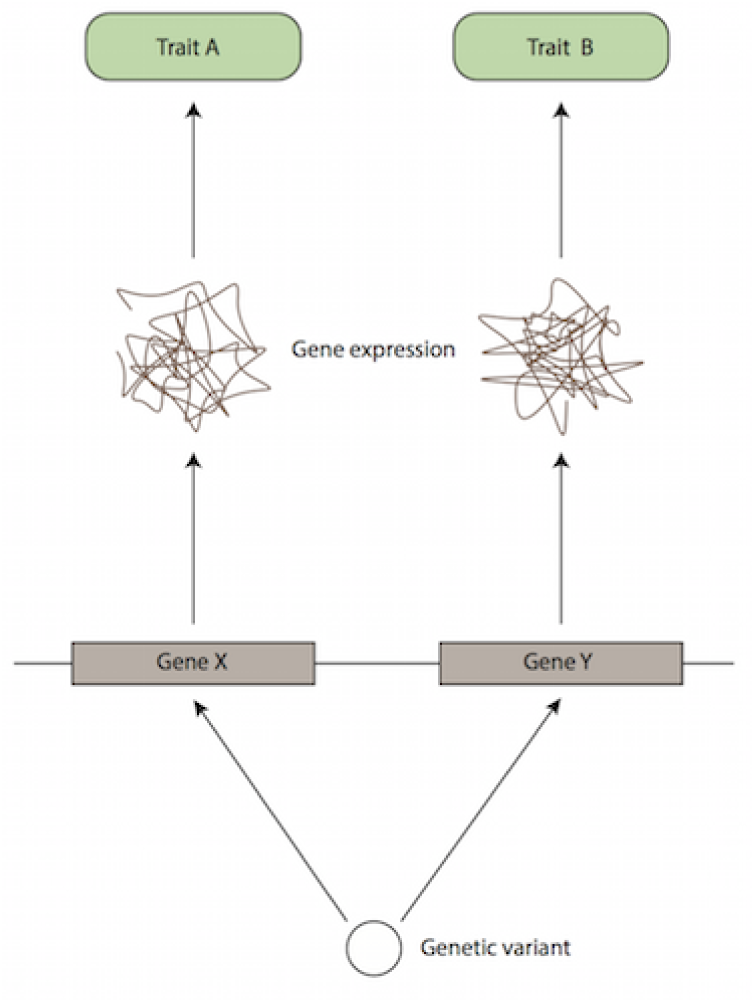
Adapted from Figure 3 in Hu et al (2016).

**Figure 3:**
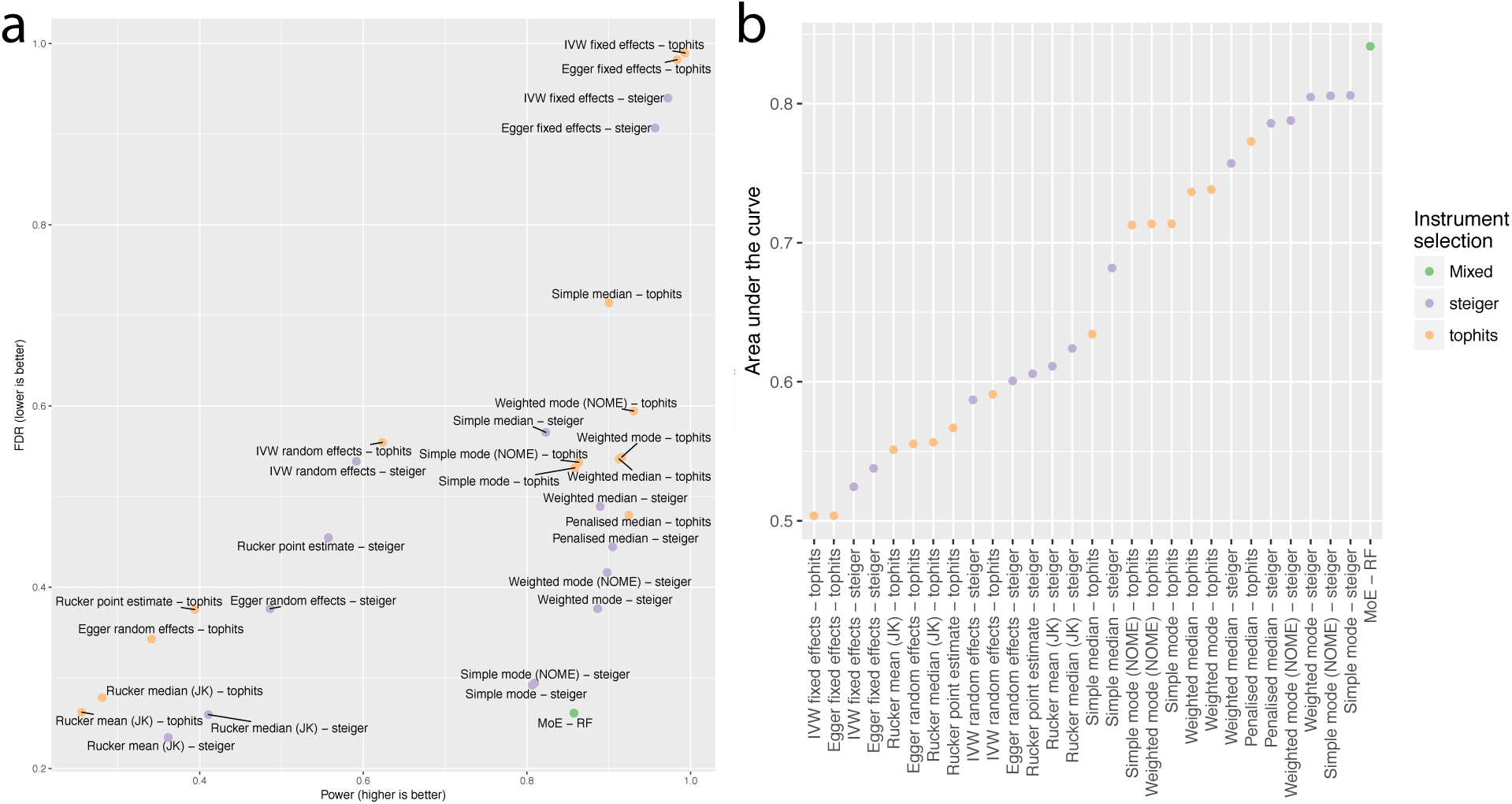
Performance of MoE against all other MR strategies. a) The power for non-null datasets is plotted against the FDR for null datasets for each of the 28 strategies, plus MoE. No single MR strategy achieved nominal FDR for these simulations. b) Calculating the area under the ROC curve from the values in (a) we plotted the performance in order from lowest to highest. Under the assumption of pervasive horizontal pleiotropy, the MoE approach is likely most effective than any other single MR strategy.

## Methods

### GWAS summary data and their use in 2SMR

The use of summary data in two-sample MR is described in detail elsewhere (12,27). A brief outline of the procedure is as follows. First, genetic instruments for the exposure trait need to be identified - those SNPs with *p* < 10^-8^ are retained in order to ensure that the first assumption of MR (that the instrument associates with the exposure) is generally satisfied. We collect their effect sizes, standard errors and effect alleles for the association with the exposure trait. These can be obtained from published GWAS summary data, either from from curated lists of established GWAS associations (10) or study websites comprising complete summary data (all SNP effects across the genome regardless of strength of association). Next, the effects of those SNPs on the outcome need to be obtained, typically necessitating access to complete summary data because these SNPs will not usually reach genome-wide significance for the outcome trait (and are therefore usually absent in curated catalogues).

We use the term *summary-set* to refer to the minimum data required to perform 2SMR analysis - four columns comprising the SNP-exposure and SNP-outcome effects and their standard errors, with each rows corresponding to a SNP that is used as an instrument for the exposure.

In its simplest implementation, the regression of the SNP-exposure effect sizes against the SNP-outcome effect sizes, with greater weight afforded to those SNPs with smaller SNP-outcome standard errors, provides the estimate of the causal effect of the exposure on the outcome. This is known as the IVW method.

Given summary data for a large number of traits, it is straightforward to exhaustively analyse the causal relationships of every trait against every other trait for which there are sufficient summary data available. Supplementary table 1 provides a list of all traits that have available GWAS summary data that were used in these analyses.

### MR methods and their assumptions

In this paper we consider three main classes of MR estimation. Full details for each approach have been described previously. A summary of the methods is given in Supplementary table 2. Though other methods also exist we have limited the analyses to the following 14 methods for simplicity.

#### Mean-based methods

Here we consider four nested models (6). The inverse variance weighted (IVW) fixed effects meta-analysis approach assumes that variants exhibit no horizontal pleiotropy. IVW random effects meta-analysis relaxes the horizontal pleiotropy assumption, allowing it to be present but balanced - such that it only leads to increased heterogeneity around the regression line without affecting the slope (and therefore not introducing bias). Fixed effects Egger regression (4) relaxes the horizontal pleiotropy assumption further by allowing a non-zero intercept which essentially allows overall horizontal pleiotropy to be directional, where its total effect influences the outcome in a specific direction. Random effects Egger regression further allows heterogeneity around the slope having accounted for overall directional horizontal pleiotropy (6), as long as the horizontal pleiotropy effects are not correlated with the SNP-exposure effects (also known as the INSIDE assumption)(4).

The Rucker framework (28), adapted to MR (6) uses estimates of heterogeneity in the IVW and Egger frameworks to navigate between these nested models. A jackknife approach (random selection with replacement of the complete set of instruments) can be used to obtain a sampling distribution for the model estimate amongst these four variations. Using 1000 rounds of jackknife estimates, we can obtain a final estimate using the mean or the median of the distribution. We only use the jackknife approach for MR analyses where there are 15 or more instruments, in order to avoid saturating the possible number of instrument combinations.

The four nested models (IVW fixed effects, IVW random effects, Egger fixed effects, Egger random effects) plus the three Rucker estimates (point estimate, mean of the jackknife, median of the jackknife) provide seven mean-based estimators.

#### Median-based methods

An alternative approach is to take the median effect of all available instruments (5,29). This has the advantage that only half the instruments need to be valid, and the estimate will remain unbiased. The weighted median estimate develops the approach further to allow stronger instruments to contribute more towards the estimate can be obtained by weighting the contribution of each instrument by the inverse of its variance. The penalised weighted median estimator introduces a further weight to the instruments, penalising any instrument that contributes substantially towards the heterogeneity statistic. Together, this provides three median-based estimators. Other estimation strategies not considered here, such as LASSO regression, have also been developed for the situation where at least half of the instruments are valid (30).

#### Mode-based methods

The mode-based estimator clusters the instruments into groups based on similarity of causal effects, and returns the final causal effect estimate based on the cluster that has the largest number of instruments (7). There are four implementations of this method: the simple and the weighted mode, each weighted with or without the assumption of no measurement error in the exposure estimates (NOME). The simple mode is the unweighted mode of the empirical density function of causal estimates, whereas the weighted mode is weighted by the inverse variance of the outcome effect. The bandwidth parameter was set to 1 by default.

### Instrument selection

#### Top hits

The simplest approach to selecting instruments is to take SNPs that have been declared significant in the published GWAS for the exposure. This typically involves obtaining SNPs that surpass *p* < 5 × 10^-8^, using clumping to obtain independent SNPs, and then replicating in an independent sample. These results are often recorded in public GWAS catalogs. Alternatively the clumping procedure can be performed using complete summary data in MR-Base (12). Complete summary data refers to association results for all SNPs used in a GWAS i.e. not only those passing statistical thresholds for significance. We call this the “top-hits” strategy.

#### Steiger filtering

With genome-wide association studies growing ever larger, the statistical power to detect significant associations that may be influencing the trait downstream of many other pathways increases. For example, if a SNP *g*_*A*_ influences trait *A*, and trait *A* influences trait *B*, then a sufficiently powered GWAS will identify the *g*_*A*_ as being significant for trait *B* (Figure 1a). Using *g*_*A*_ as an instrument to test the causal effect of *A* on *B* is perfectly valid in some cases. But the (incorrectly hypothesised) MR analysis of trait *B* on trait *A*, for example, could erroneously result in the apparent causal effect of *B* on *A*. It is to the advantage of the researcher to exclude *g*_*A*_ from the analysis from the analysis of *B* against *A*.

**Figure 1:**
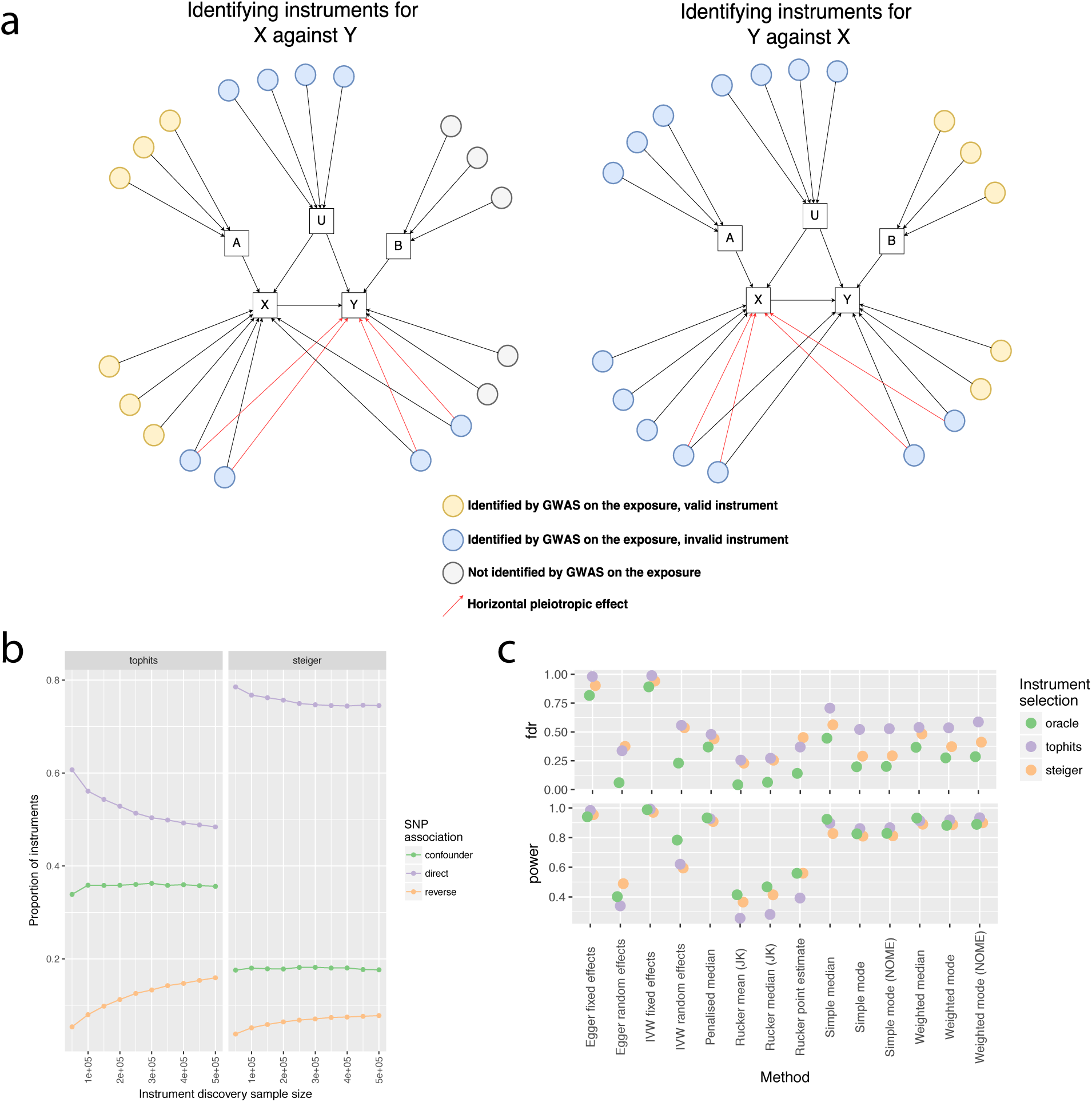
Simulations. a) Schematic of how GWAS with sufficient power can lead to the selection of instrumental variables that are invalid. We used arbitrary numbers of SNPs and confounders to simulate GWAS summary datasets. b) Any SNP that has a direct influence on the exposure, or an influence on a non-confounding intermediate variable, is considered a ‘direct’ effect. The y-axis shows the proportion of instruments selected for analysis that are either direct associations with the exposure, instruments for the outcome (reverse), or instruments for confounding traits (confounder). The proportions are compared over a range of different exposure discovery sample sizes (x-axis) and using either the top-hits approach (left) or the Steiger approach (right) for instrument selection. c) Top: The false discovery rates from null simulations for each of the 14 methods using either top-hits, Steiger filtered variants, or variants that are known to be directly associated with the exposure (oracle, note that direct effects can still exhibit horizontal pleiotropy in these simulations). Bottom: The statistical power to detect true causal associations in the non-null simulations.

An approach to inferring the causal direction between phenotypes (8) uses the following basic premise. If trait *A* causes trait *B* then

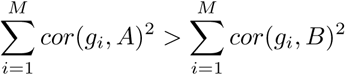

because the *cor*(*g*_*i*_*, B*)^2^ = *cor*(*A, B*)^2^*cor*(*g*_*i*_*, A*)^2^. This simple inequality will not hold in some cases, for example *cor*(*x, x*_*o*_) *< cor*(*x, y*)*cor*(*y, y*_*o*_) where *cor*(*x, x*_*o*_) and *cor*(*y, y*_*o*_) are the precision of the measurements of the *x* and *y*. Some combinations of confounding effects can also distort the *cor*(*g, x*) and *cor*(*g, y*) parameters, as has been discussed in detail previously (8). Large differences in sample sizes between the exposure and outcome GWASs may also have a practical impact on the efficacy of this approach. However, we use it here as a computationally inexpensive and approximate method to identify variants that are likely to be invalid (Figure 1a). Steiger’s Z-test of correlated correlations (31) can be used to formally test the extent to which the two correlations are statistically different.

Other methods have been developed to identify invalid instruments for the purposes of exclusion from MR analyses (9,13,32), based on the notion that outlying instruments are more likely to be a source of horizontal pleiotropy. A potential drawback of this approach is that the outlier SNPs in a summary-set might be the only reliable ones. This could arise, for example, for the *CRP* variant instrumenting C-reactive protein levels (21), or the *SLC2A9* variant influencing serum urate levels (33).

To avoid this, we primarily developed Steiger filtering to identify those instruments that are likely to be arising due to reverse cause, but it also has the potential to detect SNPs that are exhibiting horizontal pleiotropy or primarily influencing confounders. Further details are provided in the Online Methods.

#### Competitive mixture of experts

We consider the 14 MR methods described above, for which instruments can be supplied using two instrument selection strategies, leading to 28 strategies in total (Supplementary table 2). Each method assumes or performs best for a different model of pleiotropy. Our objective is to select the method most likely to be correct for a specific MR analysis by predicting the model of pleiotropy, and relating that prediction to the most appropriate model. To achieve this we use a “mixture of experts” machine learning approach (34), where each MR strategy is considered to be an ‘expert’, taking a summary-set as its input. An overview of the approach is shown in Figure 2.

**Figure 2:**
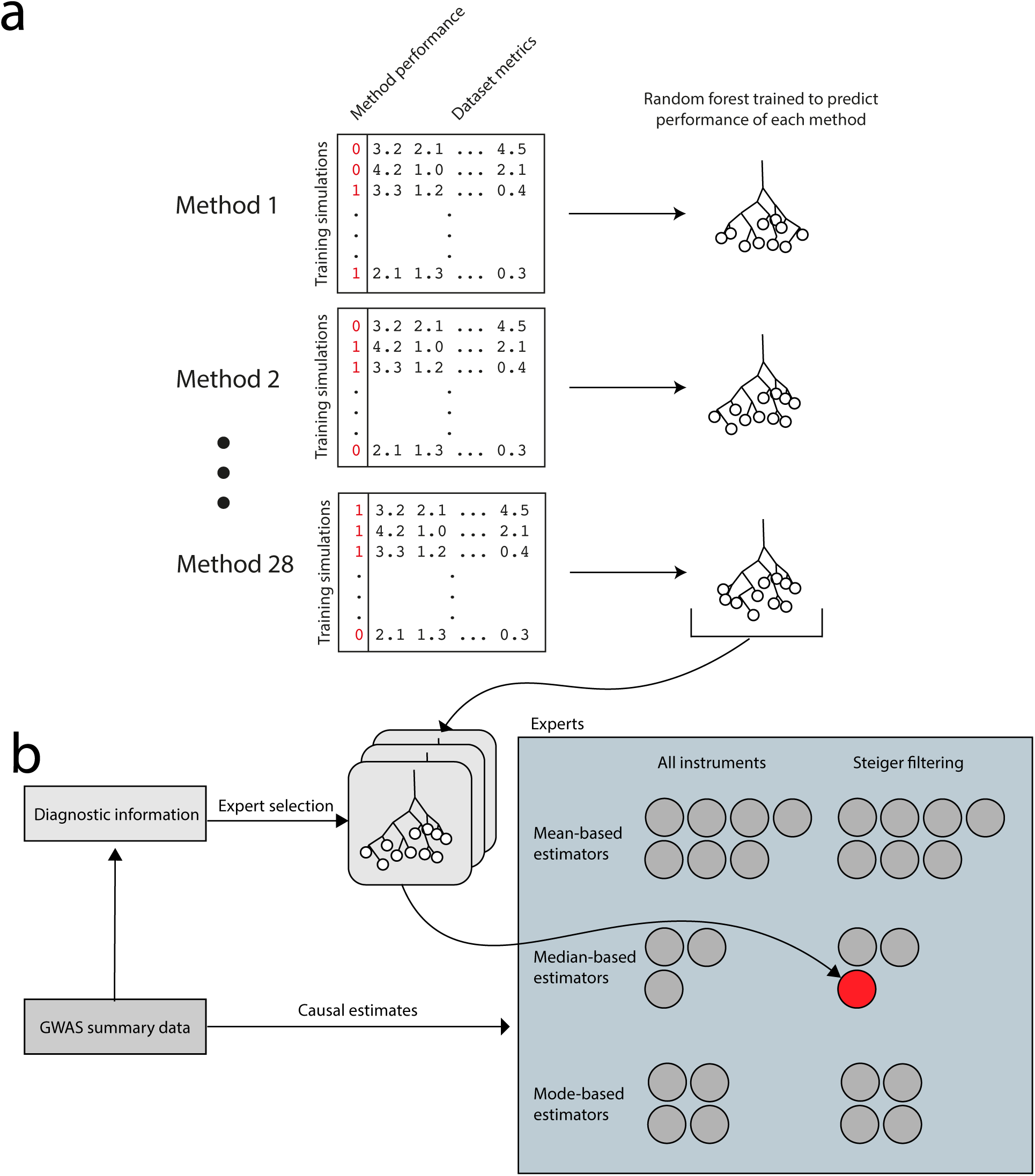
Mixture of experts. a) Training. Datasets are simulated that have either null or non-null causal relationships, and causal estimates are obtained from each of the 28 MR strategies considered in this study. In the toy datasets, the columns in black represent 53 metrics about each of the 67,000 training simulations. The columns in red are specific to each strategy, they represent how well that strategy performed in obtaining the correct answer for each of the datasets. Random forests are used to learn the parameter space of the 53 metrics in which a particular summary-set is likely to perform well. Together, this creates 28 random forest decision trees, one for each MR strategy. b) Application. For a GWAS summary dataset, our objective is to choose the strategy most likely to return the correct causal estimate. Metrics are generated from the dataset and fed into each of the 28 random forest decision trees. This provides us with 28 performance predictions. Finally, we use the strategy for which the performance prediction is highest.

The mixture of experts method seeks to divide a parameter space into subdomains, such that a particular expert is used primarily for problems that reside in a subdomain most suited to that expert. In this case we first identify characteristics of the SNP-exposure and SNP-outcome associations for which one specific MR method is most likely to yield highest statistical power for non-null associations, and lowest false discovery rates for null associations. This involves creating a ‘gating function’, whose purpose is to decide which expert to use for a specific MR analysis, given the parameter space that is occupied by that summary-set. The metrics are a collection of regression diagnostics and are described in Supplementary table 3.

The gating function needs to be trained using data for which the true causal effect is known, and to this end we generated a large number of simulated summary-sets. We trained the gating function using random forest learning algorithms that seek to identify sectors of the parameter space that are most likely to return accurate estimates for a particular expert. Figure 2a illustrates how the gating function is trained using simulated data. New summary-sets can then be applied to the trained mixture of experts (MoE) model (Figure 2b). We call this implementation MR-MoE 1.0. Full details are provided in the Online Methods. MR-MoE 1.0 is implemented in the TwoSampleMR R package available at github.com/MRCIEU/TwoSampleMR (12).

It should be noted that in the original hierarchical mixture of experts approach of (34) the gating function and the experts share the same input space and are trained simultaneously using expectation-maximisation. In our approach the gating function is defined over a separate input space consisting of the parameter space of the experts, and is trained separately in a supervised setting using simulated data. As such our MR-MoE approach fits the framework of meta-learning (35–37) where machine learning methods are applied at the ‘meta-level’ to analyse the results of machine learning experiments at the ‘base-level’, in order to be able to understand and model the capabilities of each expert and recommend which expert should be applied to a given problem. From this perspective the 28 MR strategies are the base-models, the gating function is the meta-model, and its meta-features are the parameters of the summary-sets.

### Graph database of MR estimates

The set of MR estimates obtained from this analysis, and the summary-sets used to generate them, are deposited in a Neo4j graph database. Here node representations exist for traits and SNPs; and relationship representations exist for SNP-trait associations and trait-trait MR estimates. Because each trait-trait association has up to 28 different MR estimates, for simplicity we also distill this down to a third relationship type comprising a single ‘best estimate’ for each trait-trait association using the following rules:

1. If the number of variants after Steiger filtering is greater than 5 then apply the MoE to obtain the best method. This value is chosen arbitrarily as a minimum number of variants for which the different MR methods can model horizontal pleiotropy
2. If the number of variants after Steiger filtering is less than or equal to 5 but greater than 1 then use the IVW random effects approach on the filtered set of variants
3. If there is 1 variant retained in the Steiger approach then use the Wald ratio on the remaining variant
4. If there are no variants remaining after Steiger filtering then declare no estimate of a causal association possible.

The graph can be queried directly using the cypher language at http://eve-neo4j.mrbase.org or through a basic web interface at http://eve.mrbase.org. For specific hypotheses we strongly recommend that estimates from all sensitivity analyses are scrutinised and reported.

## Results

### Steiger filtering improves reliability

As statistical power for GWAS studies improves the likelihood of a significant association being discovered for a trait that acts primarily on one of its precursors increases (Figure 1a). This presents a problem when GWAS significance is used as the sole criterion for instrument selection if the hypothesis being tested is either a) the trait causing the precursor, or b) the trait causing some other trait and the precursor causally relates to both (in which case the precursor is a confounder). We evaluated the efficacy of the Steiger filtering approach for improving instrument selection using 100,000 simulated summary-sets comprising both null and non-null causal models. For each summary-set, instruments were selected based on the top-hits strategy and the Steiger filtering strategy, and MR was performed using 14 different methods based on instruments selected from each of these strategies.

Figure 1b shows that the top-hits strategy led to over half of the instruments being primarily associated with either confounders or the outcome phenotype, not the exposure phenotype. For brevity, we refer to the latter type of invalid instruments as “reverse causal” instruments. The proportion of invalid instruments due to reverse cause increased as GWAS discovery sample size increased. Applying Steiger filtering reduced this to 25%. Consequently, the false discovery rates (FDR) for 12 of the 14 methods reduced substantially when applied using Steiger filtered instruments (Figure 1c). The true positive rates for the methods based on Steiger filtering did however reduce slightly for 10 of the 14 methods.

### Mixture of experts method selection improves over any single method

Following evidence that the Steiger filtering approach can improve on existing methods, we next hypothesised that a mixture of experts (MoE) model would be able to predict the most appropriate of the 14 MR methods and two instrument selection strategies (giving a total of 28 MR strategies) to apply to a particular summary-set based on its characteristics (Figure 2).

The ability to predict the performance for each of these methods is shown in Supplementary table 2. The prediction *R*^2^ of whether a summary-set’s status was truly null (*β*_*xy*_ = 0) or non-null (*β*_*xy*_ ≠ 0) against the method’s prediction of the summary-set’s status, ranged between 0.04 and 0.24. The method performance prediction was most effective for the Egger random effects model with Steiger filtering. The summary-set characteristics with the most importance for each of the predictors differed substantially between each summary-set, as well as the frequency for which each of the methods was selected in the testing summary-sets. The FDR of each method when chosen, compared to their averages across all summary-sets, reduced; and likewise the true positive rates of each method when chosen increased compared to their simulation-wide averages.

We compared the MoE performance in the simulations against each of the 28 strategies, testing to see if it outperformed all other single strategies. Figures 3a and 3b show that the MoE approach had the best general performance. Estimating the area under the receiver operator curve gave 0.84 for the MoE approach in terms of classifying the simulations as being null or non-null. Notably, the next best methods were median and mode based estimators using Steiger filtering. But a crucial observation is that under the assumption of widespread and diverse pleiotropic effects it is clear that all methods suffer from high false discovery rates (Figure 3a), including the MoE approach.

### Automated MR analysis of 2407 phenotypes reveals substantial horizontal pleiotropy

We applied our analysis using summary data for 2407 phenotypes, including 149 complex traits and diseases, 575 metabolites (38,39) and 1683 plasma protein levels (40). For the protein levels only GWAS significant hits were available, so they could only be evaluated as exposure phenotypes. The complex traits and diseases and metabolite levels had complete summary data available obtained via MR-Base, and thus they could be evaluated as both exposures (if they had significant instruments) and outcomes. Together, we evaluated 715681 relationships. The majority of these associations could only be evaluated using fewer than 5 SNPs, and so the Wald ratio or IVW fixed effects methods were used, but for 61029 associations the MR-MoE approach was applied.

There were 5660 associations following Bonferroni correction (*p* < 7.0 × 10^-8^). Of these 2918 were obtained from the MR-MoE analysis, while the remainder were estimated using fewer than 5 SNPs using the Wald ratio or IVW fixed effects methods.

The frequencies of the methods chosen by the MR-MoE analysis are shown in Supplementary table 3. Amongst those deemed ‘significant’, the IVW fixed effects analysis method combined with the top-hits instruments selection strategy (only valid when there is no detectable evidence of horizontal pleiotropy of any sort) was selected in only 10.4% of cases. This indicates that horizontal pleiotropy is likely to be pervasive.

### Interpretation of the putative associations in a broad scan of phenotypes requires detailed follow up

We performed a look up of associations between all traits and LDL cholesterol (as measured in the GLGC consortium (41)), where the result had a false discovery rate of 0.05 using the MoE approach. This returned 287 putative associations, amongst which 111 involved LDL cholesterol influencing other traits and 176 involved other traits influencing LDL cholesterol. A large proportion of these traits were metabolites, which are dominated by lipid fractions. Filtering to exclude the metabolomic studies ((38,39)) returned 27 associations with 23 traits (Figure 4a). There was a strong association with coronary heart disease (0.41 log(OR) per SD, 0.31-0.52, *p* = 2.7 × 10^-12^). Here the IVW random effects method was chosen after Steiger filtering, indicating the presence of horizontal pleiotropy being present amongst the instruments.

**Figure 4:**
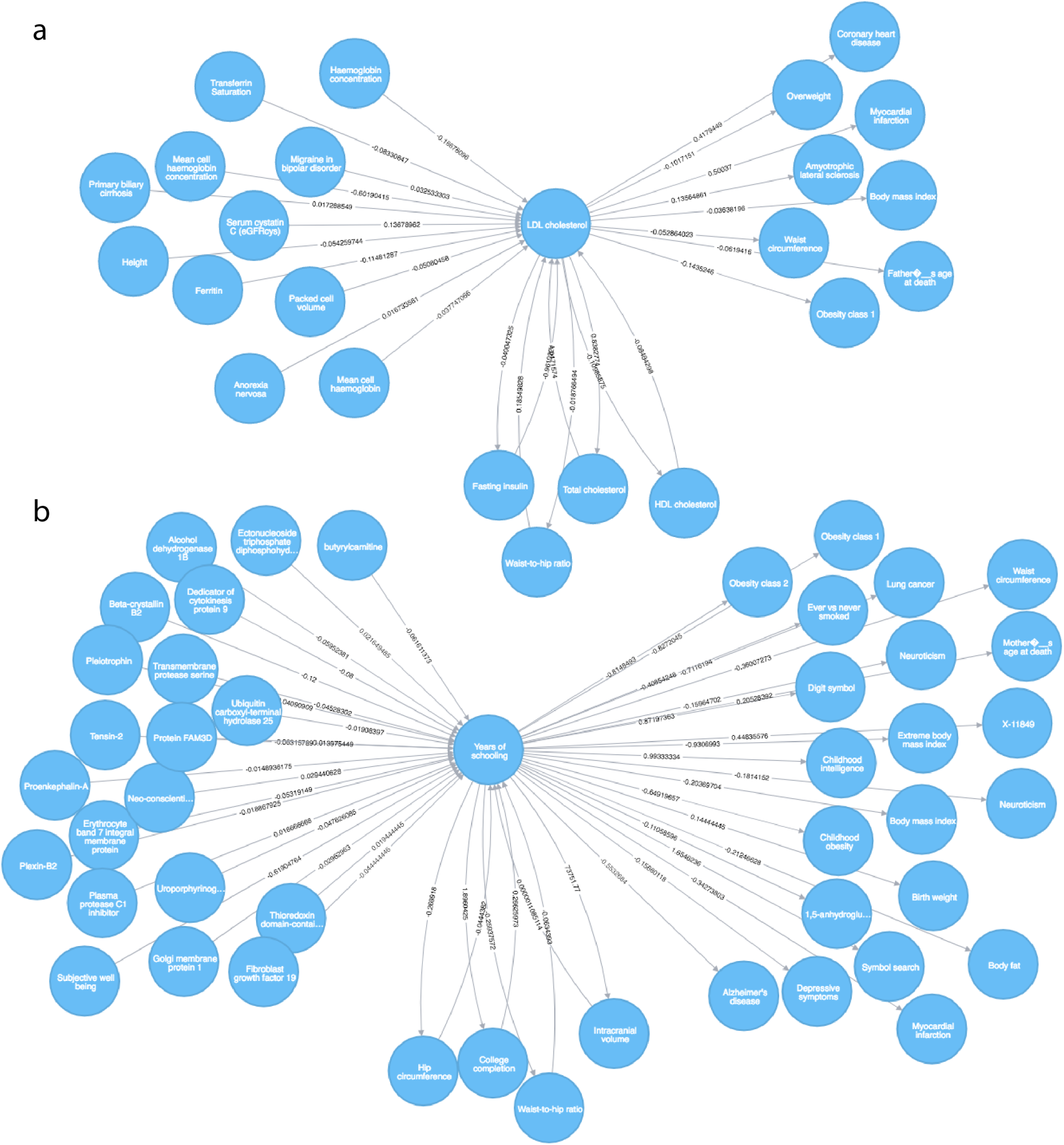
Lookup of causal associations involving a) LDL cholesterol and b) Years of Schooling. In a) the nodes are filtered to not include those obtained from the metabolomic studies (38,39). The arrows denote causal direction and the values on the arrows denote the causal effect estimate. Only those relationships are shown for which the *FDR* < 0.05.

We also performed a look up of associations that causally influence years of schooling (42), or that years of schooling itself causally influences. Using a false-discovery rate of 0.05, 45 traits were returned as having some direction of causality with years of schooling (Figure 4b). Several of these putative relationships appear to be plausible, for example the influence of years of schooling on lower risk of Alzheimer’s disease, which has recently been reported (43). There is also a protective effect of more years of schooling on lower risk of myocardial infarction, which has been supported by detailed follow up analyses (44).

However, it is clear that there are limitations in relying on MR as a panacea for causal inference. For example, bi-directional causal relationships with college completion exist because of the similarity in trait definitions; and there are several traits (e.g. childhood intelligence and birth weight) which appear to be causally downstream of years of schooling despite this being temporally impossible. Though explanations could be conjured for this association, for example parents’ schooling influences childhood intelligence, or the genetic instruments are shared across the two traits due to a third unmeasured trait, an intuitive interpretation of causality cannot be applied because the genetic instruments used proxy the liability to achieve some level of educational attainment, and not necessarily the experience of having attained a particular level of edication.

The facility to perform these look ups has been automated in the MR of ‘everything vs everything’ (MR-EvE) web application, http://eve.mrbase.org.

## Discussion

Motivated by a recent flurry of activity in MR methodology development, we have devised a machine learning approach, MR-MoE, that seeks to predict the performance of each MR method, selecting the one which is most likely to maximise power whilst minimising FDR for a specific summary-set. The mixture of experts approach that we present here is trained using random forest decision trees applied to extensive and diverse simulations, and we demonstrate that it makes substantial performance improvements over any other single method in the presence or absence of extensive horizontal pleiotropy. As such, our method contributes to the field of meta-learning but also to the rapidly growing field of automatic model selection and hyper-parameter optimization in machine learning (45). We applied the method to the curated set of GWAS summary data present in MR-Base (12), generating several million MR estimates. MR-MoE selected a method that indicated some pattern of horizontal pleiotropy for almost 90% of cases, reinforcing the notion that it is the rule rather than the exception.

The trend of increasing GWAS sample sizes continues, but while the opportunity that this affords MR to be furnished with more instruments is typically welcomed, here we have demonstrated that it is invalid instruments, exhibiting horizontal pleiotropy or reverse causation, that may be more likely to be identified than valid ones. Strategising how MR is to be used in practice, therefore, must consider some presence of horizontal pleiotropy to be the rule rather than the exception. Instrumental variables in MR are typically chosen blind, with GWAS significance being the only criterion. We illustrated how confounders and reverse causal associations can easily lead to invalid instrument selection in MR and that most randomly generated patterns of horizontal pleiotropy cannot be adequately accounted for by any single MR method.

Methods have been introduced recently that seek to improve MR estimates through invalid instrument detection and removal (9,13,32). These attack the problem from the perspective that variants which have a relatively large influence on the causal estimate, or provide substantially different estimates from the other variants, are likely to be exhibiting horizontal pleiotropy, and their removal from the analysis may be warranted. A potential drawback of this approach is that the outlier SNPs in a summary-set might be the only reliable ones. This could arise, for example, for the *CRP* variant instrumenting C-reactive protein levels (21), because CRP itself is downstream of many cellular functions and most GWAS signals will not be specific for the trait. Steiger filtering has attempted to sidestep this issue in removing invalid instruments, seeking to detect those instruments which have associations with the outcome that are too large to be consistent with the exposure causing the outcome. We acknowledge that unmeasured confounding and measurement error can introduce problems with Steiger filtering (8), but note that our simulations indicate that all individual MR methods improve in performance following its application.

In addition to horizontal pleiotropy, there are many other limitations that prevent MR from being a panacea for causal inference. Many of the associations are biologically impossible, for example where early life phenotypes appear to be influenced by later stage phenotypes. Such associations can be explained statistically because the instruments proxy the liability for that particular phenotype, not that phenotype’s observed manifestation. Though these associations can be informative, their interpretations as causal relationships are far from clear. Similarly, often disease traits appear to causally relate to other phenotypes, but GWAS of disease is typically performed on the liability scale, hence the causal estimate reflects not the presence or absence of disease, but the underlying risk of disease. Again, interpretation of such associations can be problematic due to the instruments proxying disease liability and not diseae outcomes.

A large proportion of the associations were estimated using only a single instrumental variable, a circumstance in which cause, reverse cause or horizontal pleiotropy cannot be immediately distinguished. Though methods are emerging to attempt to delineate between models of reverse cause, pleiotropy and multiple causal variants in the same region (46,47), separating vertical from horizontal pleiotropy with a single instrument is not yet possible in the two-sample MR framework though the use of mediation-based approaches may be informative in the one-sample setting (8,48).

Other problems, in addition to horizontal pleiotropy, can also manifest. Frailty effects (49) could induce associations for late-onset traits, and often detailed modelling for how this can influence MR is warranted (43,50). Genetic variants could strongly relate to several phenotypes, for example SNPs that influence LDL cholesterol are also likely to influence other lipid fractions, making it difficult to ascertain which amongst them is the true causal exposure (51). We see this issue manifesting in Figure 4, where there appear to be causal relationships between different lipid fractions. The measured feature for which genetic associations are known could relate in complicated ways to the biological entities that is truly causal (2), for example higher circulating levels of the natural antioxidant extracellular superoxide dismutase (EC-SOD) appears to causally relate to higher risk of coronary heart disease (52) because the phenotypic measure relates inversely to the EC-SOD levels in arterial walls, hence *in situ* activity is lower (2). Dynastic effects can also confound MR estimates, arising when the exposure trait in a previous generation influences the environment of the current generation. Here, the SNPs being used as instruments for the exposure will be associated with a potential confounder in the analysis (53). The associations depicted in Figure 4 involving educational attainment may be susceptible to bias arising from this mechanism, and protecting against it will likely require alternative study designs, for example by using sibling pairs.

A new problem may arise when using machine learning approaches to infer the correct model for a particular summary-set, namely, that the model selection is data driven. This is liable to contribute to elevated type 1 error rates, even though we have attempted to separate the information used for optimisation from the information used to predict performance. One advantage to generating a method that consolidates across many methods is that it can prevent cherry picking the most appealing method based on their results. The extent to which a machine-based method selector incurs higher type 1 error rates more than a human selector is not clear. Though MR-MoE does exhibit higher type 1 error rates than are desirable, they still remain amongst the lowest compared to other MR methods that do not suffer from the issue of data driven model selection.

In light of these issues, we emphasise that we do not consider the reporting of MR-MoE (or any other single MR method) to be seen as performing a Mendelian randomization study. Rather, it can be used to motivate further detailed follow up which should include sensitivity analyses (12), incorporating biological knowledge regarding instruments, and triangulation with other sources of evidence or experimental designs (14). Demanding a rigorous follow up of putative associations is also essential for avoiding issues that may arise due to cherry picking of MR results. This is especially important when a large repository of putative associations, such as MR-EvE, is made publicly available.

The field of MR is evolving. The advent of GWAS databases (10,54) and automated 2SMR (12) has trivialised the analytical aspect of investigating specific causal hypotheses. Despite the limitations described above, the construction of a causal graph of ‘everything versus everything’ does have appeal. First, though causality is not guaranteed by MR, it can still be highly informative for supporting or negating specific hypotheses. Indeed, a well-powered MR that negates a hypothesis is likely less fallible to the pitfalls described above than using MR to confirm a positive association (55). Second, it paves the way for new approaches to search for novel putative associations and improve reliability in single analyses. The potential to exploit GWAS summary data within the properties of graph databases to aid with both of these endeavours could be transformative in biological research.

## Acknowledgements

This work was supported by the UK Medical Research Council (MC_UU_12013/1, MC_UU_12013/8, MR/N501906/1); Cancer Research UK (C52724/A20138, C18281/A19169); and the SPHERE Interdisciplinary Research Collaboration funded by the UK Engineering and Physical Sciences Research Council under grant EP/K031910/1

## Online methods

### Steiger filtering

An approach to inferring the causal direction between phenotypes recently developed (8) uses the following basic premise. If trait *A* causes trait *B* then *cor*(*g*_*i*_*, A*)^2^ *> cor*(*g*_*i*_*, B*)^2^ because the *cor*(*g*_*i*_*, B*)^2^ = *cor*(*A, B*)^2^*cor*(*g*_*i*_*, A*)^2^. This will be true under most circumstances, but some parameters of measurement error or unmeasured confounding can lead to the inequality reversing direction, as has been explored previously (8).

The Steiger test is applied to each variant in turn and we exclude any *g*_*A*_ for which *cor*(*g*_*i*_*, A*)^2^ *> cor*(*g*_*i*_*, B*)^2^, indicating that it is unlikely to primarily associate with *B* relative to *A*. Similarly, for SNPs that influence confounders of *A* and *B* or exhibit horizontal pleiotropy, the difference in *cor*(*g*_*i*_*, A*)^2^ and *cor*(*g*_*i*_*, B*)^2^ will be reduced, increasing the likelihood of the SNP being excluded because the Steiger Z-test is less likely to be significant. Hence we also exclude any *g*_*A*_ for which the *cor*(*g, x*) and *cor*(*g, y*) are not significantly different at an arbitrary threshold of *p* > 0.05.

To estimate *cor*(*g, x*)^2^, if *x* is continuous we obtain the F-statistic from the reported p-value and sample size and then 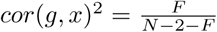. If *x* is binary then we estimate the variance of the underlying liability explained by the SNP, 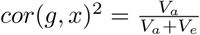. Here, *V*_*e*_ = π^2^/3, and *V*_*a*_ = 2 β^2^ *f*(1 – *f*), where β is the log odds ratio and *f* is the allele frequency of the SNP in the population (56). *f* can be estimated using the allele frequency of the SNP in an ascertained sample by deriving the 2 × 2 contingency table from the odds ratio *e^β^*, allele frequency in the ascertained sample *f*_*cc*_, and number of cases *N*_1_ and controls *N*_0_. A Z-score to test the hypothesis that *cor*(*g, x*) and *cor*(*g, y*) are significantly different can be generated using the Steiger test for correlated correlations.

### Mixture of experts implementation for MR

The mixture of experts approach seeks to use simulations to learn the circumstances in which each of the 28 MR strategies is likely to perform most accurately. Having trained the model on simulations, new summary-sets can be applied to obtain causal estimates from the strategy that is predicted to be the most reliable for that summary-set.

### Training and testing simulations

The MoE is trained using summary-sets generated from simulations (Figure 2a). Each *summary-set* can be fed into any of the 28 experts to obtain MR causal effects. The simulations used to generate these summary-sets seek to cover a range of pleiotropic scenarios, including where some proportion of SNPs exhibit directional or balanced horizontal pleiotropy, or where SNPs influence confounding variables.

We simulate two individual level datasets for which there are *N*_*x*_ and *N*_*y*_ samples, and *M* SNPs, where each SNP has effect allele frequency of *f*_*m*_ ∼ *U* (0.05, 0.95). These datasets are used to obtain the SNP effects for the exposure trait *x* and the outcome trait *y*, respectively, to create summary-sets, using the following sampling criteria:

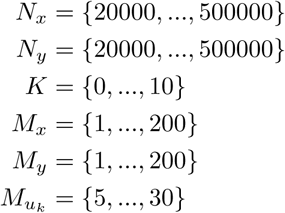

The *M* = *M*_*x*_ + *M*_*y*_ + Σ *M*_*uk*_ SNPs can influence *x* directly, *y* directly, or some number of confounders *u*_*k*_ directly. Phenotypes for *x* and *y* are constructed using

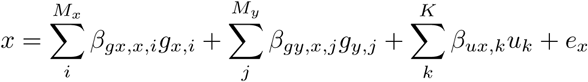

where *β*_*gx,x*_ is the vector of effects of each of the *M*_*x*_ SNPs that influence *x* primarily, *β*_*gy,x*_ is the vector of effects for the *M*_*y*_ SNPs on *x*, where the *M*_*y*_ SNPs influence *y* primarily but exhibit horizontal pleiotropic effects on *x*. We allow some proportion of these effects to be 0. *β*_*ux*_ is the vector of effects of each of the *K* confounders on *x*. Each *u*_*k*_ variable is constructed using

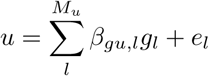

and finally *y* is constructed using

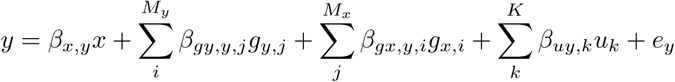

where *β*_*x,y*_ is the causal effect of *x* on *y*. We sample the distribution of direct SNP effects using

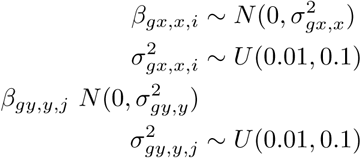

Some proportion *floor*(*M S*_*x*_*/M*) of *g*_*x*_ SNPs and *floor*(*M S*_*y*_*/M*) of *g*_*y*_ SNPs, where *s*_*x*_ and *s*_*y*_ ∼ *U* (0, 1), exhibit horizontal pleiotropy with effects sampled using

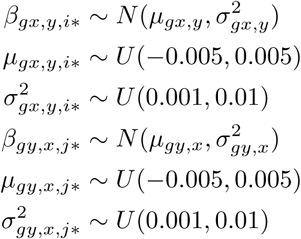

The genetic influences on each of the confounders are sampled using

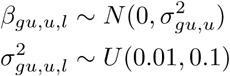

The influence of each confounder on *x* and *y* is obtained using

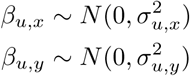

Finally, 20% of the simulations have a null effect of *β*_*x,y*_ = 0, while the other remaining 80% have a true effect sampled from

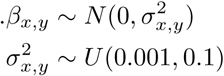

For each simulation we used linear regression to estimate the genetic effect of each SNP *M* on *x* in sample 1, and each SNP *M* on *y* in sample 2. We then perform MR analysis in both directions, mimicking GWAS by retaining SNPs that have *p* < 5*e* - 8 in sample 1 to perform MR of *x* on *y* (the true causal direction for non-null simulations), and retaining SNPs that have *p* < 5*e* - 8 in sample 2 to perform MR of *y* on *x* (the reverse causal direction for non-null simulations). We treat the summary data (effect sizes and standard errors) used for estimating *x* → *y* the summary data used for estimating *y* → *x* as two separate summary-sets. Hence, for each simulation two summary-sets are generated which are analysed to produce 28 MR estimates each. We performed 100,000 simulations using these parameters, resulting in 200,000 summary-sets.

### Optimisation function

We aim to maximise statistical power for summary-sets where *β*_*x,y*_ ≠ 0 and minimise false discovery rates for summary-sets where *β*_*x,y*_ = 0. To train random forest decision trees to predict performance for a particular method *h*(*O*_*w,d*_, **z**_*d*_) is generated where the training set of input metrics for summary-set *d* is **z**_*d*_ and the response (optimisation function) is

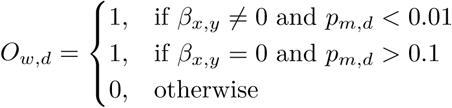

where *p*_*m,d*_ is the p-value for method *m* on summary-set *d*.

### Strategy

For each training summary-set we record a set of 53 metrics **z**_*d*_ (Supplementary table 3), and an outcome *O*_*w,d*_, which is a measure of how well that method performed for each particular simulated summary-set. For each of our 28 MR strategies, we need to create a model that predicts the performance of the strategy based on metrics generated from a summary-set. To do this, for each strategy we train a random decision forest to predict that strategy’s performance using the summary-set’s metrics. The random forest approach is well suited to this problem because there are likely to be non-continuous combinations of different metrics that improve on prediction over, for example, a simple linear model that does not learn about interactions.

As a simple hypothetical example - if a summary-set exhibits a single outlier but is otherwise exhibiting no heterogeneity then the following methods could arguably perform well:

- an IVW fixed effects analysis with Steiger filtering, should the Steiger test be able to detect the outlier
- a median based approach without Steiger filtering
- a mode based approach without Steiger filtering

deciding between these strategies requires finding, in general, which will minimise false discovery rates and maximise true positive rates for that particular scenario. In this example the IVW with Steiger filtering would likely be the clear winner because of its superior statistical power. Countless other scenarios could arise. For example if 100% of instruments are invalid but the InSIDE assumptio is met then the MR-Egger is likely to be most effective; if 40% of instruments are invalid then a median-based approach is likely most effective; and if 80% of instruments are invalid then a mode-based approach is likely most effective.

Having generated random forest decision trees for each of the 28 strategies using 133,000 of the simulations, we then applied them to the remaining 67,000 summary-sets to predict which method would have the highest performance for each of the remaining summary-sets. Finally we compare the performance of the method selected by the MoE against all remaining strategies. The default settings for the randomForest package in R (57) were used to train the models. MR-MoE 1.0 is implemented in the TwoSampleMR R package available at github.com/MRCIEU/TwoSampleMR (12).

## References

[1]. Davey Smith G, Ebrahim S. 'Mendelian randomization': can genetic epidemiology contribute to understanding environmental determinants of disease? International Journal of Epidemiology [Internet]. 2003 Feb;32(1):1–22. Available from: http://www.ije.oxfordjournals.org/cgi/doi/10.1093/ije/dyg070

[2]. Davey Smith G, Hemani G. Mendelian randomization: genetic anchors for causal inference in epidemiological studies. Human molecular genetics. 2014 Jul;23(R1):R89–R98.

[3]. Holmes MV, Ala-Korpela M, Smith GD. Mendelian randomization in cardiometabolic disease: challenges in evaluating causality. Nature Reviews Cardiology [Internet]. 2017 Jun; Available from: http://www.ncbi.nlm.nih.gov/pubmed/28569269 http://www.nature.com/doifinder/10.1038/nrcardio.2017.78

[4]. Bowden J, Davey Smith G, Burgess S. Mendelian randomization with invalid instruments: effiect estimation and bias detection through Egger regression. International Journal of Epidemiology. 2015;44(2):512–25.

[5]. Bowden J, Davey Smith G, Haycock PC, Burgess S. Consistent Estimation in Mendelian Randomization with Some Invalid Instruments Using a Weighted Median Estimator. Genetic Epidemiology [Internet]. 2016 May;40(4):304–14. Available from: http://www.ncbi.nlm.nih.gov/pubmed/27061298 http://www.pubmedcentral.nih.gov/articlerender.fcgi?artid=PMC4849733 http://doi.wiley.com/10.1002/gepi.21965

[6]. Bowden J, Del Greco M F, Minelli C, Davey Smith G, Sheehan N, Thompson J. A framework for the investigation of pleiotropy in two-sample summary data Mendelian randomization. Statistics in Medicine [Internet]. 2017; Available from: http://doi.wiley.com/10.1002/sim.7221

[7]. Hartwig FP, Smith GD, Bowden J. Robust inference in summary data Mendelian randomization via the zero modal pleiotropy assumption. International Journal of Epidemiology [Internet]. 2017;dyx102:1–14. Available from: https://oup.silverchair-8.

[8]. Hemani G, Tilling K, Davey Smith G. Orienting The Causal Relationship Between Imprecisely Measured Traits Using Genetic Instruments. bioRxiv [Internet]. 2017; Available from: http://biorxiv.org/content/early/2017/03/15/117101

[9]. Verbanck M, Chen C-Y, Neale B, Do R. Widespread pleiotropy confounds causal relationships between complex traits and diseases inferred from Mendelian randomization. bioRxiv [Internet]. 2017; Available from: http://www.biorxiv.org/content/early/2017/06/30/157552

[10]. Hindor? LA, Junkins HA, Hall PN, Mehta JP, Manolio TA. A Catalog of Published Genome-Wide Association Studies, Available at http://www.genome.gov/gwastudies. Accessed 12/10/2010. 2010.

[11]. Pierce BL, Burgess S. Efficient design for Mendelian randomization studies: subsample and 2-sample instrumental variable estimators. American journal of epidemiology [Internet]. 2013 Oct;178(7):1177–84. Available from: http://www.pubmedcentral.nih.gov/articlerender.fcgi?artid=3783091{\&}tool=pmcentrez{\&}rendertype=abstract

[12]. Hemani G, Zheng J, Wade KH, Laurin C, Elsworth B, Burgess S, et al. MR-Base: a platform for systematic causal inference across the phenome using billions of genetic associations. bioRxiv. 2016;10.1101/07.

[13]. Zhu Z, Zheng Z, Zhang F, Wu Y, Trzaskowski M, Maier R, et al. Causal associations between risk factors and common diseases inferred from GWAS summary data. bioRxiv [Internet]. 2017; Available from: http://www.biorxiv.org/content/early/2017/07/26/168674

[14]. Lawlor DA, Tilling K, Davey Smith G. Triangulation in aetiological epidemiology. International Journal of Epidemiology [Internet]. 2017 Jan;19(R1):dyw314. Available from: https://academic.oup.com/ije/article-lookup/doi/10.1093/ije/dyw314

[15]. Wright S. Evolution and the Genetics of Populations. In: Evolution and the genetics of populations. Chicago: University of Chicago Press; 1968.

[16]. Wagner GP, Zhang J. The pleiotropic structure of the genotype–phenotype map: the evolvability of complex organisms. Nature Reviews Genetics [Internet]. 2011 Mar;12(3):204–13. Available from: http://www.nature.com/doifinder/10.1038/nrg2949

[17]. Hill WG, Zhang X-S. Assessing pleiotropy and its evolutionary consequences: pleiotropy is not necessarily limited, nor need it hinder the evolution of complexity. Nature Reviews Genetics [Internet]. 2012 Feb;13(4):296. Available from: http://www.ncbi.nlm.nih.gov/pubmed/22349131

[18]. Hartwig FP, Davies NM, Hemani G, Davey Smith G. Two-sample Mendelian randomization: avoiding the downsides of a powerful, widely applicable but potentially fallible technique. International Journal of Epidemiology [Internet]. 2016 Dec;45(6):1717–26. Available from: https://academic.oup.com/ije/article-lookup/doi/10.1093/ije/dyx028

[19]. Xu L, Lin SL, Schooling CM. A Mendelian randomization study of the effiect of calcium on coronary artery disease, myocardial infarction and their risk factors. Scientific Reports [Internet]. 2017 Feb;7:42691. Available from: http://www.ncbi.nlm.nih.gov/pubmed/28195141 http://www.pubmedcentral.nih.gov/articlerender.fcgi?artid=PMC5307362 http://www.nature.com/articles/srep42691

[20]. Larsson SC, Burgess S, Michaëlsson K, TB H, WH C, Y P. Association of Genetic Variants Related to Serum Calcium Levels With Coronary Artery Disease and Myocardial Infarction. JAMA [Internet]. 2017 Jul;318(4):371. Available from: http://jama.jamanetwork.com/article.aspx?doi=10.1001/jama.2017.8981

[21]. Prins BP, Abbasi A, Wong A, Vaez A, Nolte I, Franceschini N, et al. Investigating the Causal Relationship of C-Reactive Protein with 32 Complex Somatic and Psychiatric Outcomes: A Large-Scale Cross-Consortium Mendelian Randomization Study. Hay PJ, editor. PLOS Medicine [Internet]. 2016 Jun;13(6):e1001976. Available from: http://www.ncbi.nlm.nih.gov/pubmed/27327646 http://www.pubmedcentral.nih.gov/articlerender.fcgi?artid=PMC4915710 http://dx.plos.org/10.1371/journal.pmed.1001976

[22]. Inoshita M, Numata S, Tajima A, Kinoshita M, Umehara H, Nakataki M, et al. A significant causal association between C-reactive protein levels and schizophrenia. Scientific Reports [Internet]. 2016 Sep;6(1):26105. Available from: http://www.ncbi.nlm.nih.gov/pubmed/27193331 http://www.pubmedcentral.nih.gov/articlerender.fcgi?artid=PMC4872134 http://www.nature.com/articles/srep26105

[23]. Hu JX, Thomas CE, Brunak S. Network biology concepts in complex disease comorbidities. Nat Rev Genet [Internet]. 2016 Oct;17(10):615–29. Available from: http://dx.doi.org/10.1038/nrg.2016.87 http://10.0.4.14/nrg.2016.87

[24]. Solovie? N, Cotsapas C, Lee PH, Purcell SM, Smoller JW. Pleiotropy in complex traits: challenges and strategies. Nat Rev Genet [Internet]. 2013 Jun;14(7):483–95. Available from: http://www.nature.com/doifinder/10.1038/nrg3461 http://dx.doi.org/10.1038/nrg3461 10.1038/nrg3461

[25]. Hodgkin J. Seven types of pleiotropy. The International journal of developmental biology [Internet]. 1998;42(3):501–5. Available from: http://www.ncbi.nlm.nih.gov/pubmed/9654038

[26]. Paaby AB, Rockman MV. The many faces of pleiotropy. Trends in Genetics [Internet]. 2013 Feb;29(2):66–73. Available from: http://www.ncbi.nlm.nih.gov/pubmed/23140989 http://www.pubmedcentral.nih.gov/articlerender.fcgi?artid=PMC3558540 http://linkinghub.elsevier.com/retrieve/pii/S0168952512001692

[27]. Burgess S, Scott RA, Timpson NJ, Davey Smith G, Thompson SG, EPIC-InterAct Consortium. Using published data in Mendelian randomization: a blueprint for efficient identification of causal risk factors. European Journal of Epidemiology [Internet]. 2015 Jul;30(7):543–52. Available from: http://www.ncbi.nlm.nih.gov/pubmed/25773750 http://www.pubmedcentral.nih.gov/articlerender.fcgi?artid=PMC4516908 http://link.springer.com/10.1007/s10654-015-0011-z

[28]. Rucker G, Schwarzer G, Carpenter JR, Binder H, Schumacher M. Treatment-effiect estimates adjusted for small-study efiects via a limit meta-analysis. Biostatistics [Internet]. 2011 Jan;12(1):122–42. Available from: http://www.ncbi.nlm.nih.gov/pubmed/20656692 https://academic.oup.com/biostatistics/article-lookup/doi/10.1093/biostatistics/kxq046

[29]. Han C. Detecting invalid instruments using L 1-GMM. Economics Letters. 2008;101(3):285–7.

[30]. Kang H, Zhang A, Cai TT, Small DS. Instrumental Variables Estimation With Some Invalid Instruments and its Application to Mendelian Randomization. Journal of the American Statistical Association. 2016;111(513):132–44.

[31]. Steiger JH. Tests for comparing elements of a correlation matrix. Psychological Bulletin. 1980;87(2):245–51.

[32]. Corbin LJ, Richmond RC, Wade KH, Burgess S, Bowden J, Smith GD, et al. BMI as a Modifiable Risk Factor for Type 2 Diabetes: Reffining and Understanding Causal Estimates Using Mendelian Randomization. Diabetes [Internet]. 2016 Oct;65(10):3002–7. Available from: http://www.ncbi.nlm.nih.gov/pubmed/27402723 http://www.pubmedcentral.nih.gov/articlerender.fcgi?artid=PMC5279886

[33]. White J, Sofat R, Hemani G, Shah T, Engmann J, Dale C, et al. Plasma urate concentration and risk of coronary heart disease: a Mendelian randomisation analysis. The Lancet Diabetes & Endocrinology. 2016;

[34]. Jordan MI, Jacobs RA. Hierarchical Mixtures of Experts and the EM Algorithm. Neural Computation [Internet]. 1994 Mar;6(2):181–214. Available from: http://www.mitpressjournals.org/doi/10.1162/neco.1994.6.2.181

[35]. Brazdil P, Carrier CG, Soares C, Vilalta R. Metalearning: Applications to data mining. Springer Science & Business Media; 2008.

[36]. Smith-Miles KA. Cross-disciplinary perspectives on meta-learning for algorithm selection. ACM Computing Surveys (CSUR). 2009;41(1):6.

[37]. Lemke C, Budka M, Gabrys B. Metalearning: a survey of trends and technologies. Artiflcial intelligence review. 2015;44(1):117–30.

[38]. Shin S-Y, Fauman EB, Petersen A-K, Krumsiek J, Santos R, Huang J, et al. An atlas of genetic infiuences on human blood metabolites. Nature genetics [Internet]. 2014 Jun;46(6):543–50. Available from: http://dx.doi.org/10.1038/ng.2982

[39]. Kettunen J, Demirkan A, Wurtz P, Draisma HHM, Haller T, Rawal R, et al. Genome-wide study for circulating metabolites identifies 62 loci and reveals novel systemic efiects of LPA. Nat Commun [Internet]. 2016 Mar;7. Available from: http://dx.doi.org/10.1038/ncomms11122 http://10.0.4.14/ncomms11122

[40]. Sun BB, Maranville JC, Peters JE, Stacey D, Staley JR, Blackshaw J, et al. Consequences Of Natural Perturbations In The Human Plasma Proteome. bioRxiv [Internet]. 2017; Available from: http://biorxiv.org/content/early/2017/05/05/134551

[41]. Willer CJ, Schmidt EM, Sengupta S, Peloso GM, Gustafsson S, Kanoni S, et al. Discovery and reffinement of loci associated with lipid levels. Nature Genetics [Internet]. 2013;45(11):1274–83. Available from: http://www.nature.com/doifinder/10.1038/ng.2797

[42]. Okbay A, Beauchamp JP, Fontana MA, Lee JJ, Pers TH, Rietveld CA, et al. Genome-wide association study identifies 74 loci associated with educational attainment. Nature [Internet]. 2016 May;533(7604):539–42. Available from: http://www.ncbi.nlm.nih.gov/pubmed/27225129 http://www.pubmedcentral.nih.gov/articlerender.fcgi?artid=PMC4883595 http://www.nature.com/doifinder/10.1038/nature17671

[43]. Anderson E, Wade KH, Hemani G, Bowden J, Korologou-Linden R, Davey Smith G, et al. The Causal effiect Of Educational Attainment On Alzheimer's Disease: A Two-Sample Mendelian Randomization Study. bioRxiv [Internet]. 2017; Available from: http://www.biorxiv.org/content/early/2017/04/17/127993

[44]. Tillmann T, Vaucher J, Okbay A, Pikhart H, Peasey A, Kubinova R, et al. Education and coronary heart disease: a Mendelian randomization study. bioRxiv [Internet]. 2017; Available from: http://www.biorxiv.org/content/early/2017/02/06/106237

[45]. Kottho? L, Thornton C, Hoos HH, Hutter F, Leyton-Brown K. Auto-WEKA 2.0: Automatic model selection and hyperparameter optimization in WEKA. Journal of Machine Learning Research. 2016;17:1–5.

[46]. Zhu Z, Zhang F, Hu H, Bakshi A, Robinson MR, Powell JE, et al. Integration of summary data from GWAS and eQTL studies predicts complex trait gene targets. Nature Genetics [Internet]. 2016 Mar;48(5):481–7. Available from: http://www.nature.com/doifinder/10.1038/ng.3538

[47]. Richardson TG, Zheng J, Davey Smith G, Timpson NJ, Gaunt TR, Relton CL, et al. Causal epigenome-wide association study identifies CpG sites that infiuence cardiovascular disease risk. bioRxiv [Internet]. 2017; Available from: http://biorxiv.org/content/early/2017/04/29/132019

[48]. Wang L, Michoel T. Efficient And Accurate Causal Inference With Hidden Confounders From Genome-Transcriptome Variation Data. bioRxiv [Internet]. 2017; Available from: http://www.biorxiv.org/content/early/2017/04/19/128496

[49]. Aalen OO, Valberg M, Grotmol T, Tretli S. Understanding variation in disease risk: the elusive concept of frailty. International journal of epidemiology [Internet]. 2015 Aug;44(4):1408–21. Available from: http://www.ncbi.nlm.nih.gov/pubmed/25501685 http://www.pubmedcentral.nih.gov/articlerender.fcgi?artid=PMC4588855

[50]. Noyce AJ, Kia DA, Hemani G, Nicolas A, Price TR, De Pablo-Fernandez E, et al. Estimating the causal infiuence of body mass index on risk of Parkinson disease: A Mendelian randomisation study. Brayne C, editor. PLOS Medicine [Internet]. 2017 Jun;14(6):e1002314. Available from: http://dx.plos.org/10.1371/journal.pmed.1002314

[51]. Burgess S, Freitag DF, Khan H, Gorman DN, Thompson SG. Using multivariable Mendelian randomization to disentangle the causal efiects of lipid fractions. PLOS one [Internet]. 2014 Jan;9(10):e108891. Available from: http://journals.plos.org/plosone/article?id=10.1371/journal.pone.0108891

[52]. Juul K, Tybjaerg-Hansen A, Marklund S, Heegaard NHH, Steffiensen R, Sillesen H, et al. Genetically Reduced Antioxidative Protection and Increased Ischemic Heart Disease Risk: The Copenhagen City Heart Study. Circulation [Internet]. 2003 Dec;109(1):59–65. Available from: http://www.ncbi.nlm.nih.gov/pubmed/14662715 http://circ.ahajournals.org/cgi/doi/10.1161/01.CIR.0000105720.28086.6C

[53]. Hinke S von, Davey Smith G, Lawlor D, Propper C, Windmeijer F. Genetic markers as instrumental variables. J Health Econ. 2016;45:131–48.

[54]. Staley JR, Blackshaw J, Kamat MA, Ellis S, Surendran P, Sun BB, et al. PhenoScanner: a database of human genotype–phenotype associations. Bioinformatics [Internet]. 2016 Oct;32(20):3207–9. Available from: http://www.ncbi.nlm.nih.gov/pubmed/27318201 http://www.pubmedcentral.nih.gov/articlerender.fcgi?artid=PMC5048068 https://academic.oup.com/bioinformatics/article-lookup/doi/10.1093/bioinformatics/btw373

[55]. Vanderweele TJ, Tchetgen Tchetgen EJ, Cornelis M, Kraft P. Methodological challenges in mendelian randomization. Epidemiology (Cambridge, Mass) [Internet]. 2014 May;25(3):427–35. Available from: http://www.ncbi.nlm.nih.gov/pubmed/24681576

[56]. Lee SH, Wray NR. Novel genetic analysis for case-control genome-wide association studies: quantification of power and genomic prediction accuracy. PLOS One. 2013;8(8):e71494.

[57]. Liaw A, Wiener M. Classiffication and Regression by randomForest. R News [Internet]. 2002;2(3):18–22. Available from: http://cran.r-project.org/doc/Rnews/

